# Growth of Lahontan Cutthroat Trout from multiple sources reintroduced into Sagehen Creek, CA

**DOI:** 10.1101/2021.03.18.436085

**Authors:** Jonathan E. Stead, Virginia L. Boucher, Peter B. Moyle, Andrew L. Rypel

## Abstract

Lahontan Cutthroat Trout *Oncorhynchus clarkii henshawi* have experienced massive declines in their native range and are now a threatened species under the US Endangered Species Act. A key management goal for this species is re-establishing extirpated populations using translocations and conservation hatcheries. In California USA, two broodstocks (Pilot Peak and Independence Lake) are available for translocation, in addition to potential wild sources. Yet suitability of these sources for re-introduction in different ecosystem types and regions remains an open and important topic. We conducted growth experiments using Lahontan Cutthroat Trout stocked into Sagehen Creek, CA USA. Experiments evaluated both available broodstocks and a smaller sample of wild fish translocated from a nearby creek. Fish from the Independence Lake source had significantly higher growth in weight and length compared to the other sources. Further, Independence Lake fish were the only stock that gained weight on average over the duration of the experiment. Our experiments suggest fish from the Independence Lake brood stock may be useful for re-introduction efforts into small montane headwater streams in California.

## Introduction

Lahontan Cutthroat Trout (*Oncorhynchus clarkii henshawi*) are endemic to the Lahontan Basin of northeast California, north Nevada, and south Oregon (Behnke 1972; Behnke 1992; Moyle 2002; Peacock et al. 2018). Cutthroat trout as a parent species was historically the dominant native inland trout in streams of the western US and appears to descend from “Truckee Trout” *Oncorhynchus belli* 10 mya (Smith and Stearley 2018). The Lahontan subspecies is noted for its ability to thrive in alkaline waters as they occur in the Great Basin physiographic province (Moyle 2002). Over the last 150 years, Lahontan Cutthroat Trout have vanished from the majority of their distribution due to massive stream ecosystem alterations (Griffith 1988; Schroeter 1998; Dunham et al. 2002; Moyle et al. 2011; Peacock and Dochtermann 2012). Originally listed as Endangered under the US Endangered Species Act in 1970, Lahontan Cutthroat Trout were re-classified to Threatened in 1975, in part to facilitate increased management. Yet even with substantial efforts in recent years, most populations face a high risk of extinction over the next century due to presence of non-native trout species (Peacock and Kirchoff 2004), degraded and fragmented habitats (Dunham et al. 1997; Novinger and Rahel 2003) and climate change (Moyle et al. 2017; Muhlfeld et al. 2018).

Lea (1968) originally recognized the importance of Independence Lake Lahontan Cutthroat Trout as one of few remaining self-reproducing and genetically “pure” or unhybridized populations of the subspecies. Further, Lea (1968) described how Independence Lake fish were historically connected to Sagehen Creek and likely shared individuals among sub-populations. Genomic distinctiveness of subpopulations, including Independence Lake, were later confirmed (Peacock et al. 2018), and several other distinct population segments of Lahontan Cutthroat Trout have since been identified (Peacock and Kirchoff 2004; Peacock et al. 2017). However, the sole native population remaining in the Truckee River basin occurs in Independence Lake proper, Nevada County, California, and its tributary, Independence Creek (Gerstung 1988; Peacock et al. 2017). Other populations in the basin are a product of re-establishment efforts, and wild populations are restricted to small headwater creeks isolated from non-native trout (Moyle 2002; Haak et al. 2010).

Translocations are the primary management tool for reversing Cutthroat Trout declines in their native range (Harig et al. 2000). Although ethical issues surrounding translocations are complex (Minteer and Collins 2010), these actions can be effective when combined with maintenance of natural habitat and re-opening of historical immigration routes (Schultz et al. 2018; Smith and Stearley 2018). In California, there are two cultivated strains (broodstocks) of Lahontan Cutthroat Trout available for translocations: Pilot Peak and Independence Lake. Yet because of uncertainty in historical distributions, and conditions over which subspecies occurred (Moyle et al. 2017), it is unclear which stock might perform best in various re-establishment efforts.

The objective of our study was to compare early-life growth of several sources of Lahontan Cutthroat Trout reintroduced into Sagehen Creek, CA. Sagehen Creek is a small headwater mountain meadow stream in the Truckee River basin which historically shared connectivity to Independence Lake (Lea 1968). While we primarily sought to evaluate performance of two available hatchery sources, we also included a limited evaluation of wild-collected and translocated fish as an additional frame of reference.

## Methods

Sagehen Creek is located on the eastern slope of the Sierra Nevada Mountains approximately 12 km north of Truckee, Nevada County, CA (Figure 1) within the Sagehen Experimental Forest, where recent large-scale disturbance has been minimal. Sagehen Creek is a small, spring-fed stream originating from mountain snowmelt (~2,530 m elevation) that meanders through 10 km of forest and mountain meadow before reaching Stampede Reservoir at 1,780 m elevation. Flows in Sagehen Creek are seasonally dynamic (Seegrist and Gard 1972); average discharge (1956-2005) = 0.35 m^3^ s^−1^, September base discharge = 0.06−0.08 m^3^ s^−1^, and peak discharge (in winter or spring) is typically ≥ two orders of magnitude. Average wetted stream width in the study area during August 2006 was 3.7 m (±0.1 SE).

**Figure 1.**
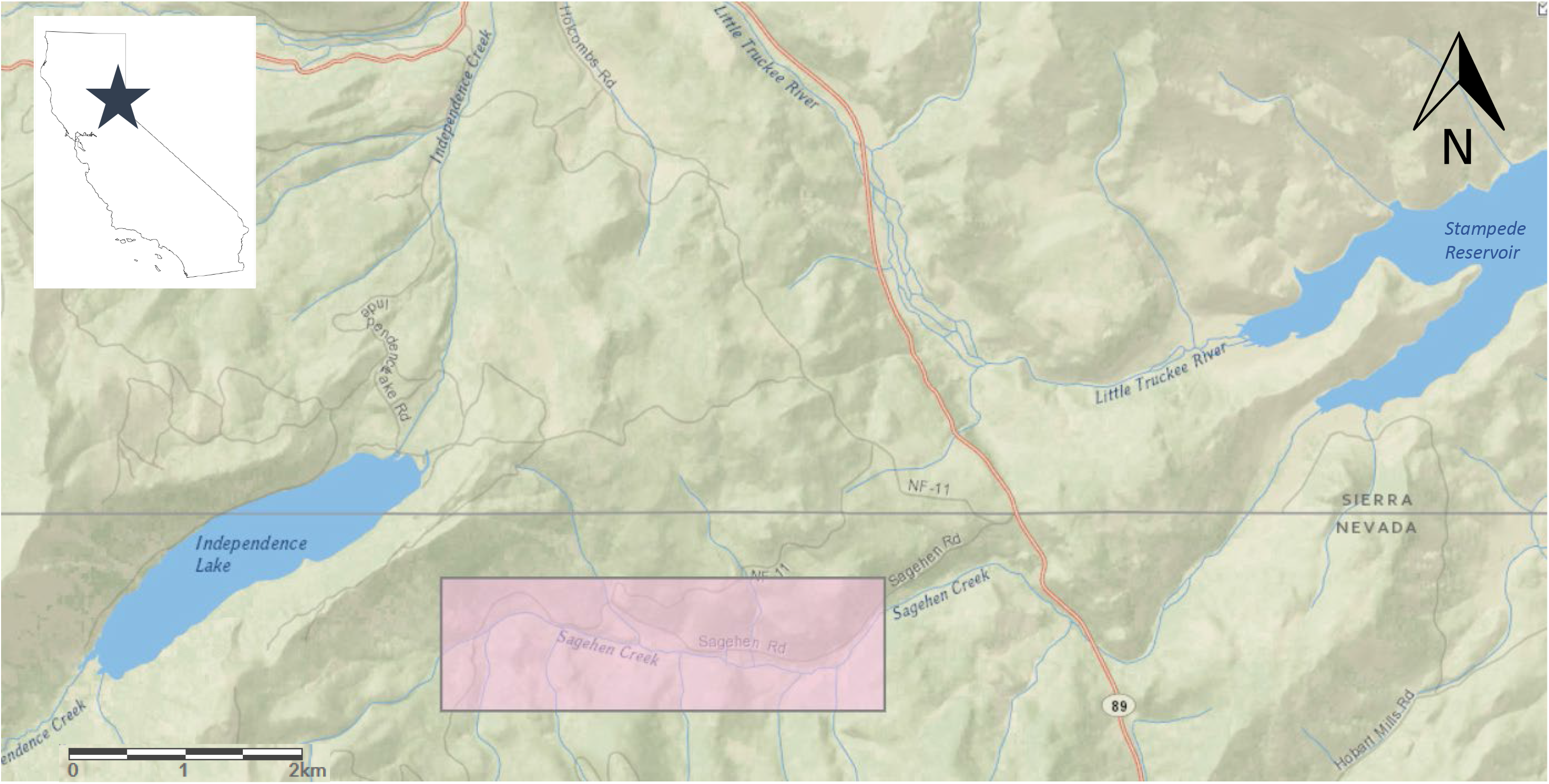
Location of Sagehen Creek, CA including study reach used in Lahontan Cutthroat Trout growth experiments, highlighted in pink.

Lahontan Cutthroat Trout disappeared from Sagehen Creek ca. 1900 coincident with extensive logging and grazing activities in the watershed. The reach where our experiment was conducted (1,950 m elevation) now supports an abundance of invertebrates, native Paiute Sculpin (*Cottus beldingii*), and naturalized populations of Brook Trout (*Salvelinus fontinalis*), Brown Trout (*Salmo trutta*), and Rainbow Trout (*Oncorhynchus mykiss*).

Two hatchery sources of Lahontan Cutthroat Trout were available for re-establishment: Pilot Peak and Independence Lake. Both sources derive from the Truckee River basin. Discovered by fish biologists in 1977 in the Pilot Peak drainage in Utah (outside the Lahontan basin), Pilot Peak fish are presumed pure Pyramid Lake stock (Hickman and Behnke 1979; Behnke 1992). The Independence Lake source derives from fish presumably isolated in the lake. Fish used in the study were originally collected by the California Department of Fish and Game from Independence Lake and planted into Heenan Lake, Alpine County, California in 1975 (Somer 2000). Reproductively mature fish in Heenan Creek (tributary to Heenan Lake) are used as a broodstock and progeny are raised in the Hot Creek Hatchery, Mono County, California USA.

Independence Lake Lahontan Cutthroat Trout were spawned on June 1, 2005 and raised by the California Department of Fish and Game (now the California Department of Fish and Wildlife or “CDFW”) at Hot Creek Hatchery for ~13 mo (Figure 2). We used an undifferentiated sample of this cohort netted from hatchery raceways and transported fish to Sagehen Creek on July 11, 2006. Pilot Peak Lahontan Cutthroat Trout were spawned during spring 2005 by the U.S. Fish and Wildlife Service at Lahontan National Fish Hatchery in Gardnerville, Nevada. Spawning occurred from late February through April, with a peak in late March. We transferred experimental fish from hatchery raceways to net pens in June Lake, Mono County, California approximately 5 months later, on September 1, 2005, where fish reared for 10 additional months. We netted an undifferentiated sample of Pilot Peak fish from pens in June Lake and transported to Sagehen Creek on July 14, 2006. To minimize differences in body condition and to habituate fish to Sagehen Creek, both hatchery groups were held in net pens in Sagehen Creek and fed to satiation on commercial trout pellets twice daily for one month prior to experiment initiation.

**Figure 2.**
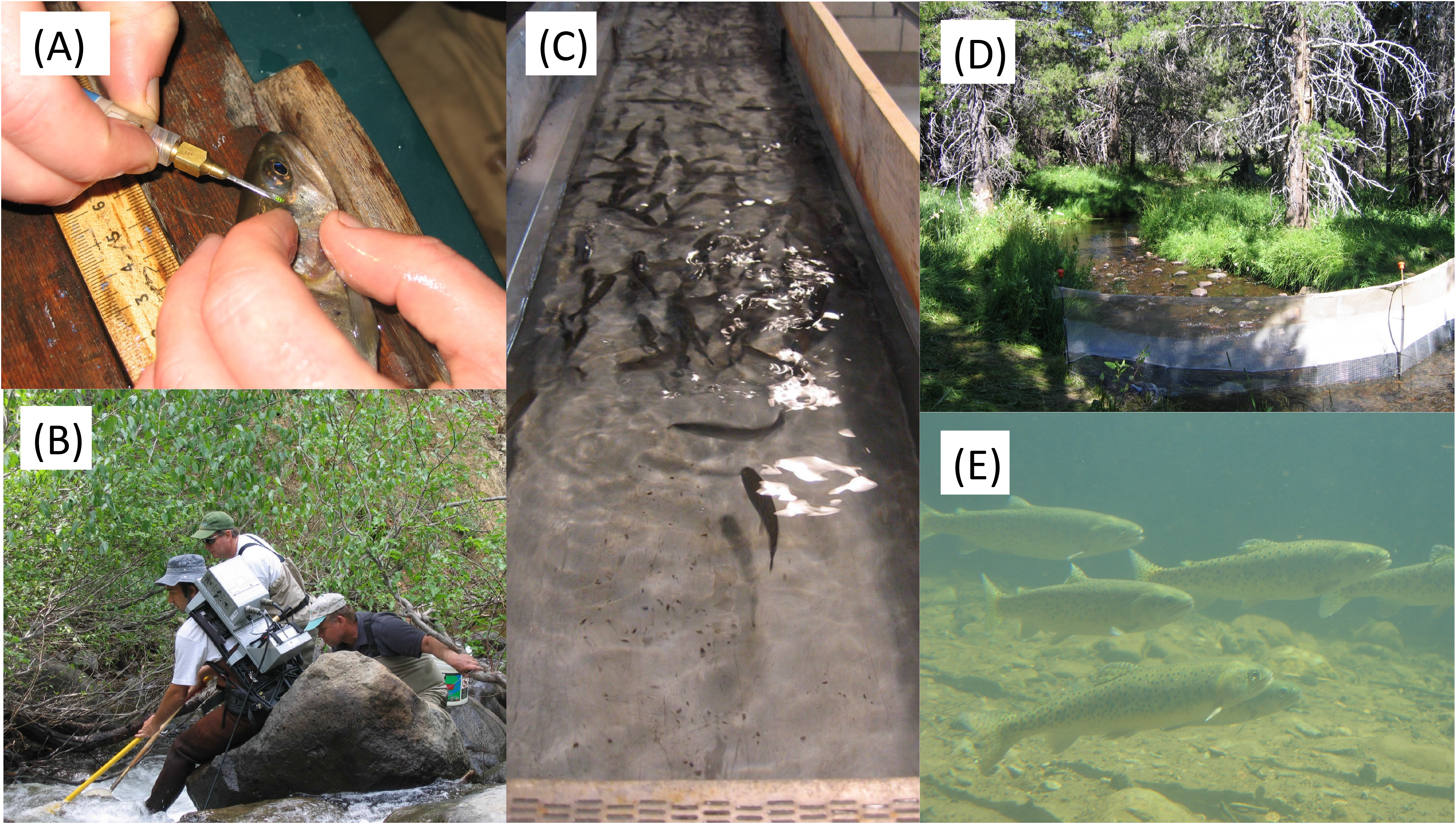
Pictures of study fish and fieldwork including (A) tagging of individual fish with visual implant alphanumeric tags; (B) sampling of fish using backpack electrofishing; (C) Lahontan cutthroat trout in the Hot Creek Hatchery; (D) Example of typical fencing and block nets used to bound one of the experimental reaches within Sagehen Creek, CA; and (E) Acclimated Lahontan cutthroat trout in Sagehen Creek, CA.

Wild Lahontan Cutthroat Trout were collected from Austin Meadow Creek (~2,075 m elevation), Nevada County, California using backpack electrofishing and hook and line sampling. This stream is a tributary of the Middle Yuba River, outside the Lahontan Basin. The population was originally stocked by the California Department of Fish and Game from Macklin Creek (also outside the Lahontan Basin) in the early 1970s. Austin Meadow fish were held in net pens for two days prior to initiating the experiment because holding wild fish in captivity prior to release may adversely affect condition and is a technique unlikely to be employed by resource agencies in future re-introduction efforts. Ultimate genetic lineages of fish used in this study were not examined; therefore we accepted the supposition that the three broodstocks represent different phenotypes and potentially genotypes of interest in future translocation efforts.

We constructed 18 temporary fish barriers enclosing nine 30 m experimental reaches over 1.7 stream km. Barriers were 1.3 cm mesh hardware cloth that enclosed the upstream and downstream sections of each 30 m reach but caused little-to-no disturbance in streamflow and drift. Each experimental reach contained a deep pool, undercut banks, woody debris and rock, cobble, boulder, and gravel substrate (Figure 2). We selected study reaches non-randomly, to control for presence of important habitat features. After completion of barriers and prior to initiation of the experiment, all non-native trout present within the study reaches were removed via multi-pass depletion using a backpack electrofisher; however all native Paiute Sculpin were returned to the stream. To the best of our knowledge, non-natives did not recolonize experimental reaches during the study.

Target stocking densities were 40 g m^−3^, representing values commonly observed from field surveys using backpack electrofishing surveys of non-native trout in Sagehen Creek (mean density = 50.3 g m^−3^; PB Moyle and VL Boucher, unpublished data), and from backpack electrofishing surveys of Lahontan Cutthroat Trout in Gance and Frazer creeks NV USA (mean density = 51.4 g m^−3^; RE Schroeter, unpublished data). Densities were adjusted downwards because: (1) Lahontan Cutthroat Trout often exist at lower densities than non-native trout (Schroeter 1998), (2) smaller Nevada streams provide greater visual buffering between fish than Sagehen Creek (RE Schroeter, personal communication), potentially resulting in higher densities (Chapman 1966), and (3) the competitive advantage of larger fish due to size dominance hierarchies (Newman 1956; Chapman 1962) may be less pronounced at low densities (Gurevitch et al. 1992). We measured habitat characteristics in each study reach including depth, volume, substrate size, canopy shade, density of large woody debris, and other habitat features. Further, we collected temperature data from the deepest part of each reach using a HOBO H8 logger over the full course of experiments (Onset Inc., Bourne MA USA, precision <0.7°C).

Stocking of Sagehen Creek began August 14, 2006 when we haphazardly netted fish from holding pens. Immediately prior to stocking, fish were sedated, fitted with 1×2.5 mm medical grade elastomer alpha-numeric visual implant tags (VI tags, Northwest Marine Technologies, Inc., Shaw Island, WA USA, Figure 2), weighed (wet weight, ± 0.05 g), and measured (standard length, mm). One VI tag was inserted into the adipose tissue behind each eye, allowing for long-term recognition of individuals. Further, recapture of enclosed fish at the conclusion of the experiment allowed some estimation of tag efficacy (Shepard et al. 1996; Turek et al. 2014), which can be higher in larger fish (Ward et al. 2015). Fish recovered ~1 h in an aerated cooler before stocking. Austin Meadow fish were stocked first, and stocking continued, alternating between hatchery sources, until addition of more fish would have resulted in a larger deviation from the target density of 40 g m^−3^ than would cessation of stocking. Stocking lasted three days total. Upon study conclusion (82 d later), fish were collected from reaches using backpack electrofishing, sedated, identified by VI tag, re-measured and -weighed, and released back to Sagehen Creek. All fish were positively identified at the end of the study based on VI tags. Mean and maximum water temperatures over all reaches ranged 6.5– 6.9°C and 0.3–15.6°C, respectively. Growth data collected during experiments are available open access in Supplementary Dataset 1.

We acknowledge growth experiments are a less common approach for managers to assess fish growth performance. For example, in many situations age and growth analyses using hard parts is preferred (Fleener 1952; Cooper 1970), or potentially tagging and recapture of wild fishes (Myrvold and Kennedy 2015; Uthe et al. 2016). Nonetheless, there is an important place within fisheries science for growth experiments, especially when evaluating translocation potential of sensitive populations in the absence of other information (Andrews et al. 2016). Translocation studies using caged fish are especially crucial to studying how Cutthroat Trout are limited by resources (Knight et al. 1999; Boss and Richardson 2002). Because Lahontan Cutthroat Trout have declined so substantially across their range, there are many basic questions about how to begin successful translocation that are best addressed using an experimental approach.

We used differences in weight and length (growth) of recaptured experimental fish to evaluate performance of fish from the three broodstocks introduced into Sagehen Creek. We initially developed weight- and length-frequency histograms for fish before and after stocking for each stock (Figure 3), and used Kolmogorov-Smirnov tests to examine whether the shape of distributions changed significantly among the two samples (periods). Comparing distributions in this manner assists with assessing potential for length-related bias in recapture probabilities between periods.

**Figure 3.**
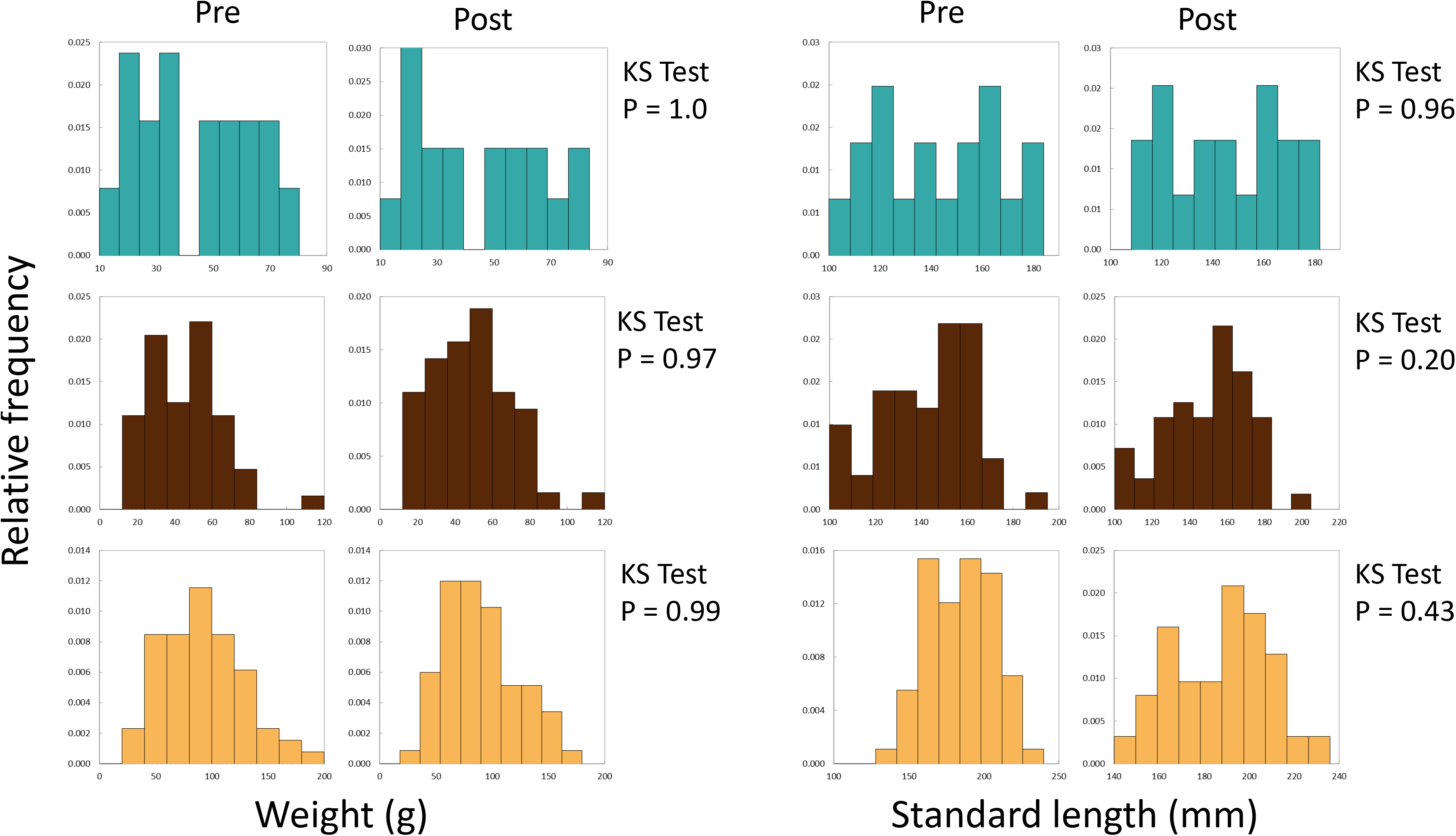
Frequency histograms for initial and final SL (top panels, open bars) and weight (bottom panels, blue bars) of all Lahontan Cutthroat Trout stocked into Sagehen Creek CA. KS Tests for SL and Weight revealed that SL and weight followed the same distributions at the beginning and end of the experiments. Black curves denote a normal distribution plotted to the data.

We compared growth rates in length and weight of Lahontan Cutthroat Trout among sources using mixed effect models fitted with restricted maximum likelihood (REML). Mixed effect models are increasingly applied to complex fisheries datasets where pseudo replication and uneven data structures are common (Daufresne et al. 2015; Rypel et al. 2016). Mixed models are also notable for their robustness when directional assumptions of distributions are violated (Schielzeth et al. 2020). Final lengths and weights for each fish were subtracted from initial lengths and weights to estimate growth for each individual fish over the course of the experiment. Normality tests revealed that weight and length growth data were mostly normal (e.g., Lilliefors Normality Tests, 5/6 Ps > 0.05). Therefore, all further analyses used raw weight and length growth data. We also note models using log-transformed or non log-transformed data yield virtually identical results. We developed two mixed effect models using growth data: one for effects of broodstock on growth in weight, and the other for growth in SL. In both models, growth in weight or SL was the dependent variable, stock was the independent variable, and reach was a random effect. Growth differences between stocks were assessed using Fisher’s Post Hoc tests. Whereas effects of habitat were controlled (to the extent possible) by creating analogous experimental stream reaches, we do not present models that analyze the effect of habitat on growth. However, we did build such models and found they differed little from above; therefore we present simple models for better accessibility. All analyses were conducted in SAS statistical software (Version 9.4, SAS Institute Inc., Cary, North Carolina, USA) and considered significant when α < 0.05.

## Results

SL and weight data followed normal distributions, and did not differ in weight or length for any stocks examined before versus after experiments (Figure 3, KS Tests, all Ps >0.20); thus there was little evidence of length-related bias in recapture probabilities. Growth rates of Lahontan Cutthroat Trout were variable under experimental conditions (Table 1, Figure 4). Across all treatment groups, negative growth in weight was common. Austin Meadow, Independence Lake and Pilot Peak fish growth ranged −7.5 – 9.8 g, −5.6– 20g, and −7.4 – 21 g, respectively. For weight, only Independence Lake showed a median value for growth that was positive (Figure 4); thus on average both Austin Meadow and Pilot Peak fish lost mass over the experiment. A mixed effect model revealed weight growth of Lahontan Cutthroat Trout varied significantly across the three broodstocks (Mixed Effect Model; −2 Res Log(Likelihood) = 827.3, Random Effects P = 0.038; Chi-Square < 0.00001). In particular, growth of Independence Lake fish was significantly faster compered to Pilot Peak (Fisher’s Test P = 0.001) and Austin Meadow (Fisher’s Test P = 0.02). Growth did not differ between Pilot Peak and Austin Meadow (Fisher’s Test P = 0.27).

**Table 1.** Numbers and change in mean length and weight of Lahontan Cutthroat Trout stocked into (n1) and collected from (n2) each reach. Independence Lake = IL, Pilot Peak = PP, and Austin Meadow = AM. ΔWeight and ΔSL data reflect the mean ±1 SE. Raw growth data from experiments can be downloaded from Supplementary Dataset 1.

| Source<br>($n_1, n_2$) | ΔWeight<br>(g) | ΔSL<br>(mm) |
| --- | --- | --- |
| Reach 1 |  |  |
| IL (7,5) | -2.0±0.5 | 7.4±1.0 |
| PP (7,7) | -5.9±2.4 | 5.4±1.5 |
| AM (2,2) | -3.0±0.9 | 4.0±0.0 |
| Reach 2 |  |  |
| IL (7,3) | -3.7±1.0 | 3.3±0.9 |
| PP (6,6) | -8.0±2.2 | 4.0±1.8 |
| AM (2,2) | -3.6±1.7 | 2.0±0.0 |
| Reach 3 |  |  |
| IL (8,5) | -1.7±0.9 | 5.4±1.5 |
| PP (8,8) | -4.3±2.6 | 4.6±1.4 |
| AM (2,2) | -2.3±0.1 | 0.5±0.5 |
| Reach 4 |  |  |
| IL (5,4) | 1.9±1.9 | 7.5±1.6 |
| PP (5,4) | -2.6±0.5 | 3.3±3.0 |
| AM (2,1) | -7.5±0.0 | -1.0±0.0 |
| Reach 5 |  |  |
| IL (7,4) | 5.2±1.1 | 6.8±2.4 |
| PP (6,6) | 2.8±2.7 | 4.0±1.8 |
| AM (3,3) | -1.0±1.1 | 2.0±2.1 |
| Reach 6 |  |  |
| IL (8,5) | 2.5±1.6 | 5.2±1.0 |
| PP (8,7) | -4.2±1.7 | 4.3±1.6 |
| AM (2,0) | no data | no data |
| Reach 7 |  |  |
| IL (10,7) | 8.4±1.9 | 8.0±0.9 |
| PP (10,9) | 5.7±2.2 | 5.9±1.5 |
| AM (2,2) | 5.4±4.5 | 5.0±3.0 |
| Reach 8 |  |  |
| IL (12,12) | 3.0±1.0 | 7.6±1.2 |
| PP (11,10) | -6.0±2.2 | -0.4±1.2 |
| AM (3,3) | 2.1±3.7 | 1.0±1.5 |
| Reach 9 |  |  |
| IL (8,7) | 2.5±0.7 | 6.6±1.1 |
| PP (8,8) | 0.7±1.1 | 3.0±1.6 |
| AM (3,3) | -0.7±2.3 | 2.3±1.5 |

**Figure 4.**
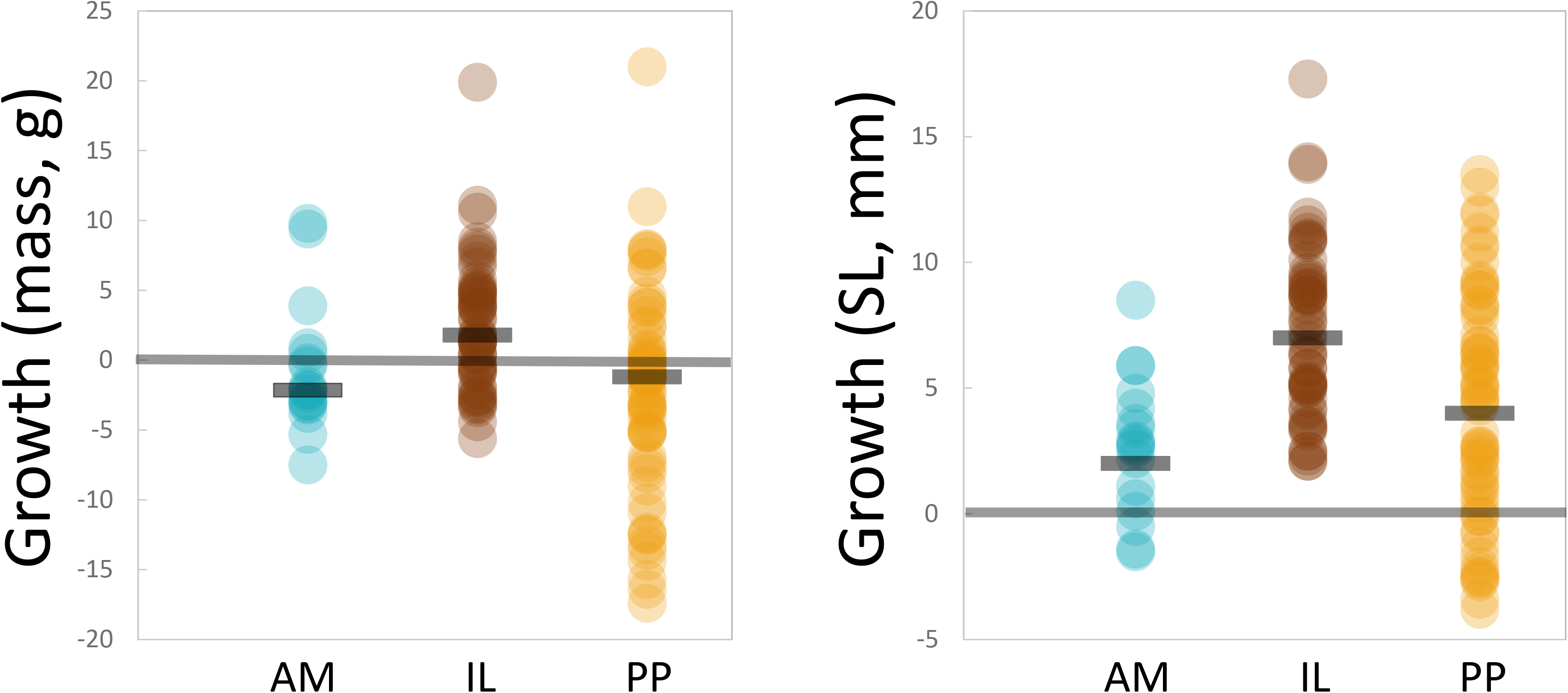
Total growth observed for individual Lahontan Cutthroat Trout over the course of the experiment. AM = Austin Meadow (orange circles), IL = Independence Lake (blue circles), PP = Pilot Peak (gray circles). Horizontal black bars = median growth for each broodstock. Yellow line denotes zero growth over the course of the experiment.

Patterns in SL growth mostly mirrored that observed for weight. However, growth in size (SL) was on average positive; all three stocks showed positive median increases in fish size (Figure 4). Austin Meadow, Independence Lake and Pilot Peak fish growth ranged −1.5–8.8 mm, 2.1–17.3 mm, and −3.8–13.5 mm, respectively. A mixed effect model showed growth in length, similar to mass, differed significantly across stocks (Mixed Effect Model; −2 Res Log(Likelihood) = 747.2, Random Effects P = 0.25; Chi-Square < 0.00001). Growth of Independence Lake fish was significantly higher compered to Austin Meadow (Fisher’s Test P < 0.0001), and Pilot Peak fish (Fisher’s Test P = 0.003). Again, growth of Austin Meadow fish did not differ significantly compared to Pilot Peak (Fisher’s Test P = 0.13).

## Discussion

Translocations are one of few tools available for management of rare and declining species (Griffith et al. 1989; George et al. 2009; Olden et al. 2011). Yet few general guidelines exist for developing successful translocation programs for freshwater fishes (but see George et al. 2009). Here we present a case study for evaluating suitability of multiple broodstocks of Lahontan Cutthroat Trout for re-introduction into a high priority Sierra Nevada stream. In our study, Independence Lake Lahontan Cutthroat Trout gained more weight and grew more in length on average compared to other evaluated stocks. In the case of weight, only Independence Lake fish showed positive growth on average under experimental conditions. We note that some individuals from all three stocks showed positive and negative growth. Furthermore, reach was a significant random effect in both models, indicating local habitat conditions are important. Independence Lake brood stock may be useful for re-introduction efforts into small montane headwater streams in California like Sagehen Creek.

Our results parallel those from similar studies in the Sierra Nevada. In a small Sierra Nevada stream, Reimers (1963) documented a high weight loss and lack of significant growth in length for Rainbow Trout (*Oncorhynchus mykiss*) during the first summer and fall over five years of stocking experiments involving multiple strains. Furthermore, initial weight loss was strongly associated with high over-winter mortality of fish, especially in late winter. Pilot Peak source Lahontan Cutthroat Trout responded to re-introduction with a weight loss consistent with data reported from Reimers (1963). We make no suppositions about contribution of rapid growth to long-term fitness of trout in Sagehen Creek; however vigorous growth of Independence fish suggests some individuals adjusted to novel conditions over the experiment.

One possible explanation for observed differences in experimental growth between hatchery sources is that Independence Lake fish are pre-adapted to headwater stream environments like Sagehen Creek. This broodstock originated from Independence Lake, located <3 km north of Sagehen Creek, immediately beyond a ridge forming the northern boundary of the watershed. Prior to construction of dams and reservoirs like Stampede Reservoir downstream (Erman 1973), Sagehen Creek and Independence Creek (outlet of Independence Lake) were adjacent tributaries of the Little Truckee River. While its name and successful brood establishment in Heenan Lake (SE of Loope CA) suggests Independence Lake fish are lacustrine-adapted fish, the degree to which these fish are truly lake-adapted is unclear. Independence Lake is actually an impoundment of Independence Creek that enlarged by ~6x a previously much smaller lake/pond from a volumetric capacity of 0.0037 km^3^ to 0.0216 km^3^ (Berris et al. 1998). Prior to construction of the dam at its outlet in 1939, the “lake” consisted of two smaller marshes that merged following impoundment. Originally, Lahontan Cutthroat Trout were thought to spend their entire life in only 20 m of stream (Miller 1957); however this perspective has shifted over time, especially in more permanent and lower order rivers. In one movement study of Lahontan Cutthroat Trout, average distance moved in three reaches of the Truckee River ranged 0.8–1.8 km (Alexiades et al. 2012). Further research in the Summit Lake Basin, NV demonstrated adfluvial behavior of resident Lahontan Cutthroat Trout and that this life history flexibility may lead to increased species persistence over time (Campbell et al. 2019).

In contrast, the Pilot Peak broodstock derives from Pyramid Lake. This ecosystem is a larger and completely natural lake with volumetric capacity = 26.4 km^3^ (Reuter et al. 1993). Lahontan Cutthroat Trout in Pyramid Lake were apex predators that had large body sizes (>18.6 kg) and low niche overlap with other conspecifics (Heredia and Budy 2018). Behnke (1992) speculated the large body size of historic Pyramid Lake fish had a genetic basis; a hypothesis that has been supported in a recent genetic study highlighting the uniqueness of this population (Peacock et al. 2017).

### Study limitations

This experiment represents a first attempt at evaluating short-term growth differences between two broodstocks of Lahontan Cutthroat Trout intended for use in re-establishment efforts. Although it allowed for a realistic evaluation of short-term response to re-introduction in a headwater stream, our approach also had issues. Due to effort required to create and maintain 18 temporary fish barriers on a daily basis, we were unable to include experimental controls, such as reaches stocked with only one source. Therefore, our results may best represent translocations in which multiple sources are used, or trout are already present in the receiving habitat. Constriction of fish movement may have altered growth results such that fish were unable to move to obtain food resources or were “forced” into artificially higher densities that aren’t encountered any longer in extant populations. Ultimately “cage effects” are necessary by-products of experiments, and while we attempt to account for these dynamics using random effects in statistical models, they are endemic to ecological experiments involving enclosed animals (Hairston 1989). We caution our limited experiments did not address these complicated issues and encourage interpretation in the context of our experimental design.

We were unable to include enough wild Austin Meadow fish because of small sample sizes of the source population (owing to the rareness of these fish) to draw strong conclusions related to that group. However, our results are not dissimilar from Ozer and Ashley (2013) who documented sanctuary stocks may not always be preferable to source stocks with higher heterozygosity. This study also focuses on growth as a measure of fitness (Roff 1983; Rypel 2011; Rypel 2014); however, additional information on long-term growth and the link between growth and survivorship would be helpful (Pedersen et al. 2017). For example, our results document short-term differences in growth among stocks during late summer and early fall summer; however, growth patterns observed during other times of the year, or across years, might be different. Furthermore, the role of competition and social cues within hierarchies may also be important (Dunham and Vinyard 1997; Knight et al. 1999; Akbaripasand et al. 2014). So-called self-thinning is frequently documented as an important process (Vøllestad et al. 2002; Lobón‐Cerviá and Mortensen 2006; Tatara et al. 2009), but it is unclear the extent to which these dynamics apply to Lahontan Cutthroat Trout. Finally, we note that our results in Sagehen Creek might simply be different if executed in other streams and ecosystem types. Additional studies are needed to continue building understanding of the mosaic of management actions needed to recover Lahontan Cutthroat Trout populations at scale.

### Conservation Management Implications

Use of the Pilot Peak broodstock is critical to recovery of Lahontan Cutthroat Trout populations in the large interconnected landscape of Truckee River and Pyramid Lake (Truckee River Basin Recovery Implementation Team 2003). However, Independence Lake fish may also be useful for re-introductions in certain situations such as in small headwater streams. Given the current status of Lahontan and other Cutthroat Trout populations, translocations will continue as an essential tool for maintaining existing populations, re-establishing new populations, and ensuring preservation of sufficient genetic variation for future evolutionary change (Schultz et al. 2018). Managers should also recognize negative relationships between reproductive performance of natural populations of salmonids and proportion of hatchery fish present (Chilcote et al. 2011); thus translocations of hatchery fish should be limited to areas where the sub-species is known to have already vanished so as to prevent introgression with existing wild stocks (Yamamoto et al. 2006).

We see a high value in Sagehen Creek specifically in pioneering upland re-establishment strategies for Lahontan Cutthroat Trout. Sagehen Creek has a rich history as a representative Sierra coldwater stream and fishery (Needham and Jones 1959; Seegrist and Gard 1972; Erman 1973; Gard and Flittner 1974; Decker 1989). For years, research focused on understanding interspecific and harvest dynamics of cold-water non-native trout species (Gard and Seegrist 1972; Erman 1986). Re-introduction of Lahontan Cutthroat Trout might be effective in Sagehen Creek if coupled with non-native species management. For example, installation of a weir, or a series of weirs, may assist in preventing colonization by non-native species. In addition, periodic non-native removals may be needed, perhaps following a high flow winter (or wild fire burn year with winter runoff) that would naturally drive Brook Trout populations to vulnerable levels (Seegrist and Gard 1972; Meyers et al. 2010). We note active fish management is a key to modern re-establishment and success of most any Lahontan Cutthroat Trout population.

## Supporting information

Length, Weight, and Growth Data

## Acknowledgements

Funding and field assistance was provided by U. S. Fish and Wildlife Service. Our study also benefited from extensive cooperation and assistance from the California Department of Fish and Game. We thank Cameron Zuber, the Moyle lab, Jim Plehn, Dan Ryan & family, Stephanie Mehalick, and others for excellent field assistance. Neil Willits at UC Davis provided valuable statistical support. Helpful reviews were provided by Sharon Lawler and two anonymous reviewers, and we appreciate discussions with Patrick Crain and Joseph Cech. Sagehen Creek Field Station at UC Berkeley provided generous facility use and equipment support. Collecting permits were provided by California Department of Fish and Game and a UC Davis Institutional Animal Care and Use Committee. ALR was supported by the Peter B. Moyle & California Trout Endowment for Coldwater Fish Conservation and the California Agricultural Experimental Station of the University of California Davis (grant number CA-D-WFB-2467-H).

